# Proteomic profiling of intracranial atherosclerotic plaque in the human brain

**DOI:** 10.1101/2024.02.10.579787

**Authors:** Qing Hao, Erming Wang, Ju Wang, Zhiping Wu, John F. Crary, Shivangi Sharma, Emma L. Thorn, Fanny Elahi, Bin Zhang, Junmin Peng

**Author notes:** corresponding authors: Qing Hao, MD, Ph.D., Assistant Professor, Department of Neurology, Icahn School of Medicine at Mount Sinai, 1468 Madison Avenue, New York, NY 10029, USA, (O) (F), Erming Wang, Ph.D., Sr. Scientist, Department of Genetics and Genomic Sciences, Icahn School of Medicine at Mount Sinai, 1425 Madison Avenue, New York, NY 10029, USA, (O). These authors contributed equally to the work (Qing Hao, Erming Wang).

## Abstract

**Background:** Intracranial atherosclerotic disease (ICAD) is one of the major causes of ischemic stroke and associated with high risk of stroke recurrence. There are no reliable and specific fluid biomarkers for ICAD, and little is known about the proteomic profiling of ICAD. In this study we aimed to explore the feasibility of applying proteomics technology to profile intracranial atherosclerotic plaques extracted from postmortem human brain arteries.

**Methods:** Eighteen segments (5-10mm in length) of major arteries from 10 postmortem brains were collected from the Mount Sinai Neuropathology Brain Bank. Among these segments, 5 had no evidence of atherosclerotic disease, and 13 had wall thickening or visible plaques with various degree of stenosis. Proteins were extracted from the vessel segments, quantified, and digested into peptides. Subsequently, the peptides underwent tandem mass tag (TMT) labeling, pooling, and analysis using two-dimensional liquid chromatography-tandem mass spectrometry (LC/LC-MS/MS). Protein identification and quantification were performed using the JUMP software. Differentially expressed proteins (DEPs) were defined as proteins with p.adj < 0.05 and absolute log2 (fold change) > log2 (1.2).

**Results:** A total of 7,492 unique proteins were detected, and 6,726 quantifiable proteins were retained for further analysis. Among these, 265 DEPs, spanning on 252 unique gene, were found to be associated with ICAD by comparing the arterial segments with vs those without atherosclerotic disease. The top 4 most significant DEPs include LONP1, RPS19, MRPL12 and SNU13. Among the top 50 DEPs, FADD, AIFM1 and PGK1 were associated with atherosclerotic disease or cardiovascular events in previous studies. Moreover, the previously reported proteins associated with atherosclerosis such as APCS, MMP12, CTSD were elevated in arterial segments with atherosclerotic changes. Furthermore, the up-regulation of APOE and LPL, the ICAD GWAS risk genes, was shown to be associated with the plaque severity. Finally, gene set enrichment analysis revealed the DEP signature is enriched for biological pathways such as chromatin structure, plasma lipoprotein, nucleosome, and protein-DNA complex, peroxide catabolic and metabolic processes, critical in ICAD pathology.

**Conclusions:** Direct proteomic profiling of fresh-frozen intracranial artery samples by MS-based proteomic technology is a feasible approach to identify ICAD-associated proteins, which can be potential biomarker candidates for ICAD. Further plaque proteomic study in a larger sample size is warranted to uncover mechanistic insights into ICAD and discover novel biomarkers that may help to improve diagnosis and risk stratification in ICAD.

## Introduction

Intracranial atherosclerotic disease (ICAD) and resultant stenosis (ICAS) are among the most common etiologies for ischemic stroke. It causes about 8-30% of ischemic strokes (the prevalence varies by race), carries a high risk of recurrence (up to 15% after one year), and leads to other adverse effects such as cognitive impairment^1–4^. Vascular factors such as hypertension, diabetes mellitus, age and smoking are well-established risk factors for ICAD. However, ICAD can also occur in patients with relatively younger age or those without significant risk factors. In addition, risk of stroke recurrence remains high despite maximal medical treatment, which includes antithrombotic medications and aggressive control of vascular risk factors^5–8^. Thus, there is a pressing need to identify novel biomarkers (beyond the traditional risk factors) that are associated with occurrence of ICAD and its associated detrimental outcome in order to improve early diagnosis and risk stratification, and inform more effective therapeutic targets to prevent ICAD and stroke attributed to ICAD.

As an emerging and powerful platform in the systematic study of proteins, proteomics analysis can identify proteins in a high-throughput matter and offers great promise in revealing complex pathways, identifying novel biomarkers for diagnosis and prognostication, monitoring treatment effects and discovering new treatment targets^9–12^. Particularly, mass spectrometry (MS)-based proteomics has demonstrated superiority in offering hypothesis-free, in-depth, multiplexed and highly specific measurements, which allows for unbiased novel biomarker discovery^13, 14^. Human proteomic studies in atherosclerotic disease have mainly focused on peripheral blood, and extracranial tissues that were obtained from endarterectomy or coronary bypass surgeries^15–23^. While a considerable number of novel markers have been described to be correlated with cardiovascular disease or degree of atherosclerosis, more efforts are underway to integrate these biomarkers into clinical practice^24–27^. Little is known about the role and significance of proteomics in characterizing molecular and cellular as well as biochemical mechanisms underlying ICAD. As intracranial arteries carry different histological and hemodynamic features compared to extracranial vessels, it is unclear whether the gene signatures derived from proteomic profiling of extracranial atherosclerosis could be generalized to intracranial lesions. In addition, proteomic profiling of peripheral blood may be affected by various co-existing conditions, whereas proteomics analysis of the plaque itself may reveal proteins that are more specific to ICAD, which helps to better understand the pathogenesis of ICAD and target biomarkers that are highly specific to ICAD^28^.

To our knowledge, direct proteomic analysis of intracranial atherosclerotic plaques has not been reported. In this study, we aim to explore the feasibility of studying ICAD plaque proteome by MS-based proteomic profiling on intracranial fresh-frozen vessels.

## Methods

### Tissues collection

A total of 18 segments of fresh intracranial arteries were collected from 10 brains recruited to the Mount Sinai Neuropathology Brain Bank & Research CoRE (NPBB) (**Supplementary data 1**). The mean age of the subjects at the time of death was 81.8 years and 6 and 4 were male and female, respectively. Four subjects had history of dementia, 7 had hypertension, 4 had hyperlipidemia, 2 had diabetes, 1 had chronic kidney disease and 2 had stroke. The presence and degree of the atherosclerotic stenosis were assessed visually based on the methodology modified following the previously published methodology for coronary artery disease^29^ (**Figure 1, upper panel**). Among the 18 vessels segments, 5 with no evidence of atherosclerotic disease was defined as Grade 0 (G0, **Figure 1, upper panel A**), 3 with wall thickening without visual plaque was defined as Grade 1 (G1, **Figure 1, upper panel B**), 3 with visible plaque with < 20% stenosis was defined as Grade 2 (G2, **Figure 1, upper panel C**), 3 with plaque with stenosis 20-50% was defined as Grade 3(G3, **Figure 1, upper panel D**), and 4 with plaque with stenosis 50-70% or > 70% was defined as Grade 4-5 (G4-5, **Figure 1, upper panel E and F**). The collected vessel samples were banked under −80 Celsius and were shipped to St. Jude Children’s Research Hospital Center for Proteomics and Metabolomics for proteomic analysis. The vessels were processed and MS-based proteomic analysis were performed as described before^30^ and outlined in **Figure 1, bottom panel**.

**Figure 1.**
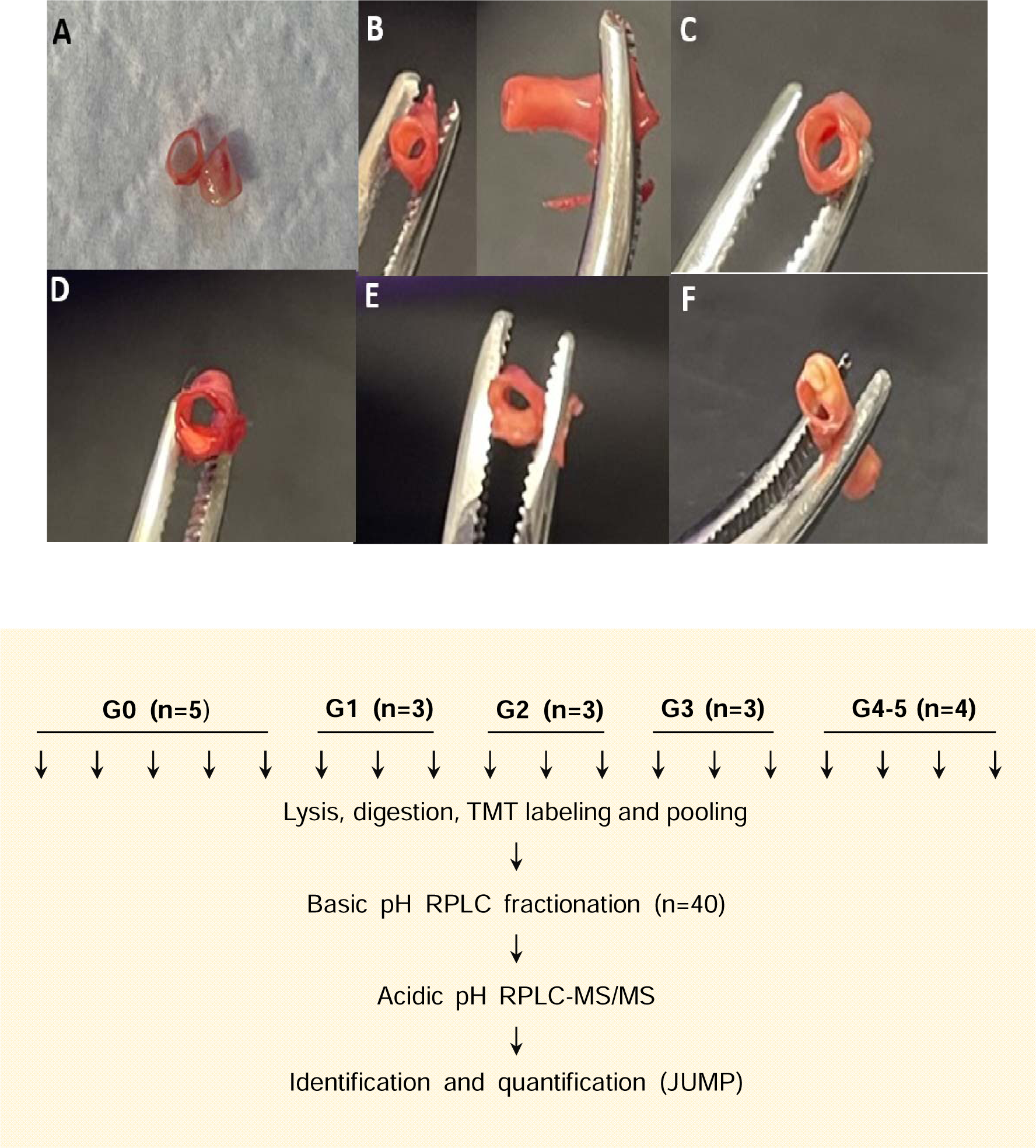
Summary of experimental materials and methods. Upper panel, visual assessment of degree of atherosclerotic disease. A, Grade 0 (G0), thin wall, no atherosclerotic disease; B, Grade 1 (G1), wall thickening, yellow appearing, no visible protrusion or stenosis; C, Grade 2 (G2), wall thickening, visible plaque protrusion, with stenosis < 20%; D, Grade 3 (G3), wall thickening, visible plaque protrusion, with stenosis 20-50%; E, Grade 4 (G4), wall thickening, visible plaque protrusion, with stenosis 50-70%; and F, Grade 5 (G5), wall thickening, visible plaque protrusion, with stenosis > 70%. Bottom panel, outlining of the proteomics profiling procedure.

### Protein extraction, digestion and Tandem-Mass-Tag (TMT) labeling

Human vessel samples underwent lysis through vortexing in a Bullet Blender in a lysis buffer (composed of 50 mM HEPES at pH 8.5, fresh 8 M urea, and 0.5% sodium deoxycholate) with the inclusion of glass beads (100 µL lysis buffer and approximately 20 µL beads for every 10 mg of tissue). Protein concentration was determined using the Pierce™ BCA protein assay kit. The protein samples were subjected to digestion with Lys-C (Wako, 1:100 w/w) at room temperature (RT) for 3 hours. Subsequently, the samples were diluted with 50 mM HEPES (pH 8.5) to reduce urea concentration to 2 M and further digested overnight with trypsin (Promega, 1:50 w/w) at RT. The digested peptides underwent reduction with fresh dithiothreitol (DTT, 1mM) for 2 hours, followed by alkylation with iodoacetamide (IAA, 10 mM) in the dark for 30 minutes. The alkylation reaction was quenched with DTT (30mM) for an additional 30 minutes. The peptides were acidified by addition of trifluoroacetic acid (TFA) to 1%, and the solution was centrifuged at 21,000 × g for 10 minutes to remove pellets. Each supernatant underwent desalting with a C18 Ultra-Micro SpinColumn (Harvard apparatus) using the standard protocol, and the peptide eluate was dried in a speedvac. Peptide samples were reconstituted in 50 mM HEPES pH 8.5 and labeled with TMTpro using a TMT/protein ratio (w/w) of 1.5:1 for 30 minutes. The labeled samples were quenched with 0.5% hydroxylamine. Equal pooling of TMTpro-labeled samples was achieved through three rounds of pooling with adjustments.

#### Basic pH LC fractionation

The combined peptide sample from the 18-plex experiment underwent fractionation through an offline basic pH reversed-phase liquid chromatography (RPLC) process utilizing an Acquity BEH C18 column (1.9 µm particle size, 3.0 mm × 15 cm, Waters). The mobile phases consisted of buffer A (10 mM ammonium formate, pH 8.0) and buffer B (90% acetonitrile, 10 mM ammonium formate, pH 8.0). The peptides were eluted over a 160-minute gradient of 15% to 42% buffer B. Fractions were collected at 0.5-minute intervals and subsequently concatenated into 40 fractions.

### Liquid Chromatography-Tandem Mass Spectrometry (LC-MS/MS) Analysis

Each fraction underwent analysis using an acidic pH reverse-phase liquid chromatography-tandem mass spectrometry (LC-MS/MS) system, employing a C18 column (CoAnn Technologies Inc., 75 μm × 20 cm with 1.7 μm C18 resin, heated at 65 °C to minimize back pressure) and a Q Exactive HF Orbitrap MS (Thermo Fisher Scientific). For peptide elution, a 78-minute gradient ranging from 13% to 58% B was employed (Here buffer A was consisted of 0.2% formic acid and 5% DMSO, and buffer B was composed of buffer A with an additional 65% acetonitrile) in an overall 95-minute run. The MS settings included MS1 scans (60,000 resolution, 460–1600 m/z scan range, 1 × 10^6^ AGC, and 50 ms maximal ion time) and 20 data-dependent MS2 scans (60,000 resolution, starting from 120 m/z, 1 × 10^5^ AGC, 110 ms maximal ion time, 1.0 m/z isolation window with 0.2 m/z offset, 32 normalized collision energy (NCE), and 10 s dynamic exclusion).

### Protein identification and quantification by JUMP software

Protein identification utilized the JUMP search engine^31^. The protein database was compiled by merging human protein sequences from Swiss-Prot, TrEMBL, and UCSC databases (83,955 entries). To assess false discovery rates (FDR), target protein sequences were concatenated with a decoy database which was generated by reversing the target protein sequences. Key parameters encompassed a 15 ppm mass tolerance for precursors, 20 ppm for product ions, full trypticity, and two maximal missed cleavages. Static modifications included Cys carbamidomethylation (+57.02146) and TMT modification of Lys and N-termini (+304.20715). Additionally, oxidation of Met (+15.99492) was considered a dynamic modification. The resulting peptide-spectrum matches (PSMs) were filtered based on mass accuracy and subsequently organized by precursor ion charge state. JUMP-based matching scores (Jscore and ΔJn) were applied with stringent cutoffs to maintain the false discovery rate below 1% for proteins. In cases where a peptide originated from multiple homologous proteins, adherence to the rule of parsimony guided the assignment to the protein with the highest PSM number. Protein quantification relied on TMT reporter ion intensities.

### Differential expression analyses

The expression (normalized read counts) of the protein groups (termed proteins) was log2-transformed. In this study, we sought to identify ICAD-associated proteins. Thus, we first classified the 18 vessels segments into two groups: Control group includes 5 segments with no evidence of atherosclerotic disease (G0); Case group includes 13 segments with atherosclerotic changes [wall thickening (G1) and visible plaque (G2, G3, and G4-5)]. We next identified the differentially expressed proteins (DEPs) in case vs control by the moderated t-test implemented in the R package limma^32^, and the nominal p values were then adjusted for multiple tests using the BH method^33^. DEPs were determined as proteins that have a false discovery rate (FDR) of less than 0.05 and an absolute fold-change > log2 (1.2). We defined these DEPs as ICAD-associated proteins. Finally, we performed gene set enrichment analysis on the DEPs over the human gene ontology database to identify their enrichment in biological processes and functional pathways that are relevant in ICAD pathology^34^. All the statistical analyses will be performed using R (version 4.2.0 or above).

## Results

Following the proteomics profiling procedure outlined in Figure 1 (Bottom panel), we identified 55,477 unique peptides, which summarized into 7,492 unique protein groups (termed proteins). Finally, we obtained a total of 6,726 unique quantifiable proteins (**Supplementary data 2a-c**). Loading bias analysis revealed that there existed no obvious loading bias (**Supplementary Figure 1**), and further principal component analysis (PCA) indicated that overall, the control group can be clearly separated from the case group (**Supplementary Figure 2**), suggesting in general good quality control (QC) was achieved in the proteome processing. Thus, the 6,726 unique quantifiable proteins constitute the protein pool (**Supplementary data 2a-c**) for further downstream analysis.

Our ultimate goal is to discover and develop biomarkers for ICAD. As the first step, we set out to identify ICAD-associated proteins, which, we reason, are a potential pool and a start point for ICAD-relevant biomarker development. The differential expression analysis is one of the ways detecting ICAD-associated proteins. Therefore, we next performed DEP analysis on these proteins and identified 265 DEPs in case vs control (**Figure 2A and 2B, Supplementary data 3**). As shown in Figure 2A and 2B, most of the DEPs were up-regulated in ICAD as compared to control. For example, the top 4 proteins including LONP1, RPS19, MRPL12 and SNU13 are up-regulated in ICAD (**Figure 2A and 2B, Supplementary data 3**). In contrast, only a limited number of proteins were down-regulated in ICAD, among which ST13 and MOB2 possessed the lowest adjusted p value (**Figure 2A and 2B, Supplementary data 3**). Further inspection of the top 50 DEPs revealed several proteins, including FADD, AIFM1, PGK1 and APOE (**Supplementary data 3**), were shown to be associated with atherosclerotic disease, cerebro- or cardiovascular events in previous studies ^35–38^.

**Figure 2.**
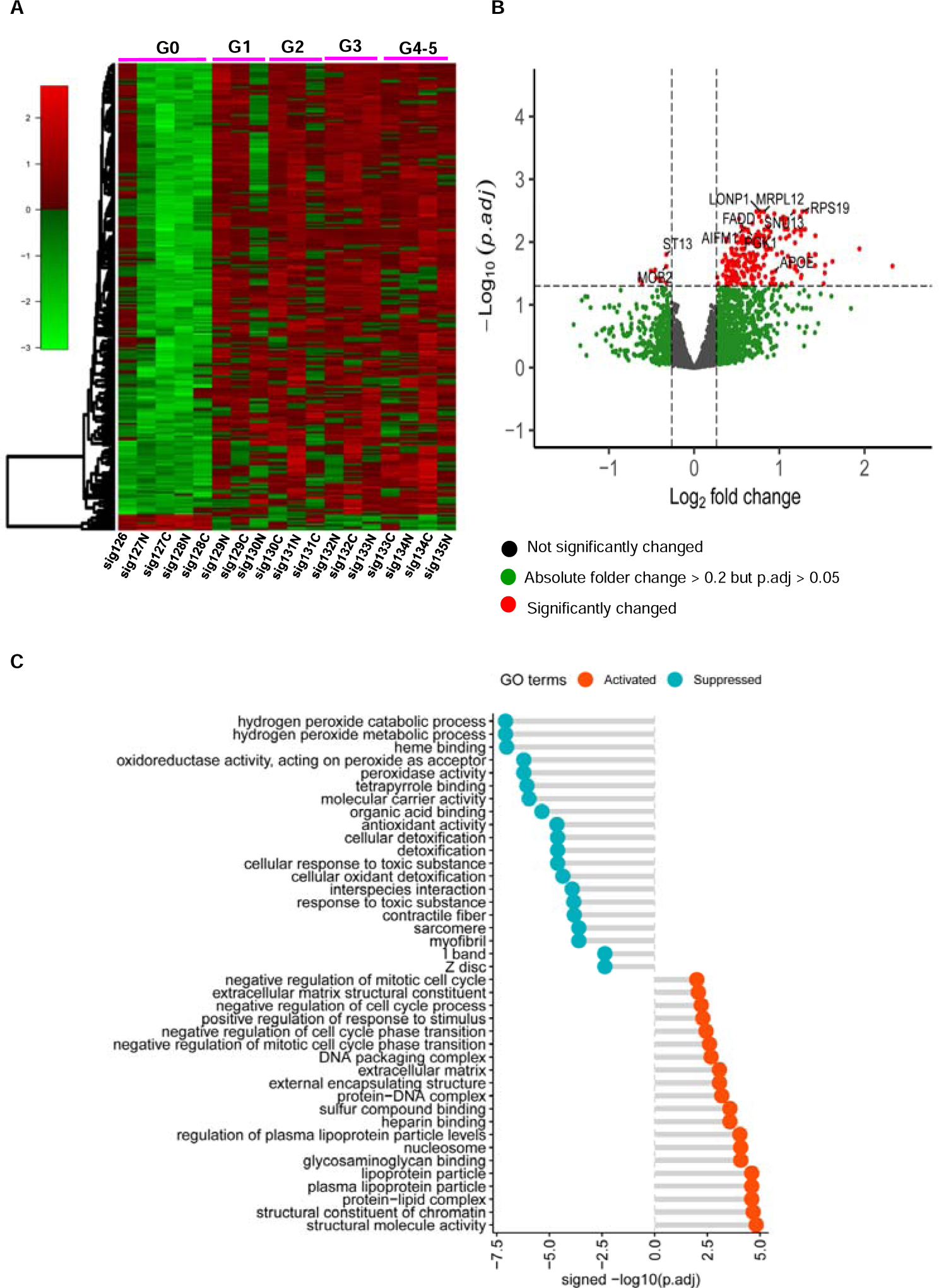
Identification and characterized of ICAD-associated differentially expressed proteins. A, Heatmap showing the expression of the DEPs across the samples. Color scheme shows row max (red) and row min (green), which represents relative expression of each protein among all the samples. B, Volcano plot for the DEP analysis. Each dot represents a protein. Highlighted are top-ranked DEPs. Since a gene could have more than 1 protein isoforms, one the isoform with the lowest p value was shown. Red dots, DEPs with p.adj < 0.05 and absolute log2 (FC) > log2 (1.2) where FC stands for fold change; green dots, proteins with p.adj ≥ 0.05 and absolute log2 (FC) > log2 (1.2); black dots, proteins with p.adj ≥ 0.05 and absolute log2 (FC) ≤ log2 (1.2). C. Gene ontology (GO) analysis. GO terms are grouped by activated (in red) or suppressed (in turquoise). Shown on the x axis are the –log10(p.adj), with the sign of minus and positive for suppressed and activated GO terms, respectively.

To gain insight into the biological pathways and functional processes the DEPs are involved in, we conducted the gene ontology (GO) analysis by gene set enrichment^34^. We detected 59 GO terms which are significantly (p value < 0.05) activated (35 GO terms) and suppressed (24 GO terms) in ICAD (**Supplementary data 4**). Figure 2C illustrated the top-ranked 20 activated and 20 suppressed GO terms, respectively. The structural constituent of chromatin, protein-lipid complex, glycosaminoglycan binding, and nucleosome are among the top activated GO terms (**Figure 2C**). In contrast, hydrogen peroxide metabolic process, peroxidase activity, antioxidant activity, and cellular detoxification are those GO terms suppressed in ICAD (**Figure 2C**). Therefore, the DEPs are involved in a range of biological pathways and functions relevant to ICAD.

Lastly we explored in detail for the functional relevance of some of the DEPs that are either relevant to atherosclerosis pathology or the ICAD GWAS risk genes^39^. Figure 3 highlighted the expression of the DEPs that were found to be associated with atherosclerosis across the 5 groups with different degrees of atherosclerotic changes. Previous research showed APCS, CTSD and MMP12 were elevated in unstable extracranial atherosclerotic plaque^40–42^. In addition, PGK1 was found to be positive in the emboli that were retrieved by thrombectomy of large vessel occlusion in acute ischemic stroke, and was associated with high LDL levels^37^. Similarly, in this study we found that the expression of these 4 proteins (APCS, CTS, MMP12 and PGK1) was increased as the degree of intracranial atherosclerotic changes worsened (Figure 3A-D). Strikingly, the expression of the ICAD GWAS risk genes, APOE and LPL^38, 39, 43^, were increased as the ICAD disease severity gets worse (Figure 3E and F).

**Fig. 3.**
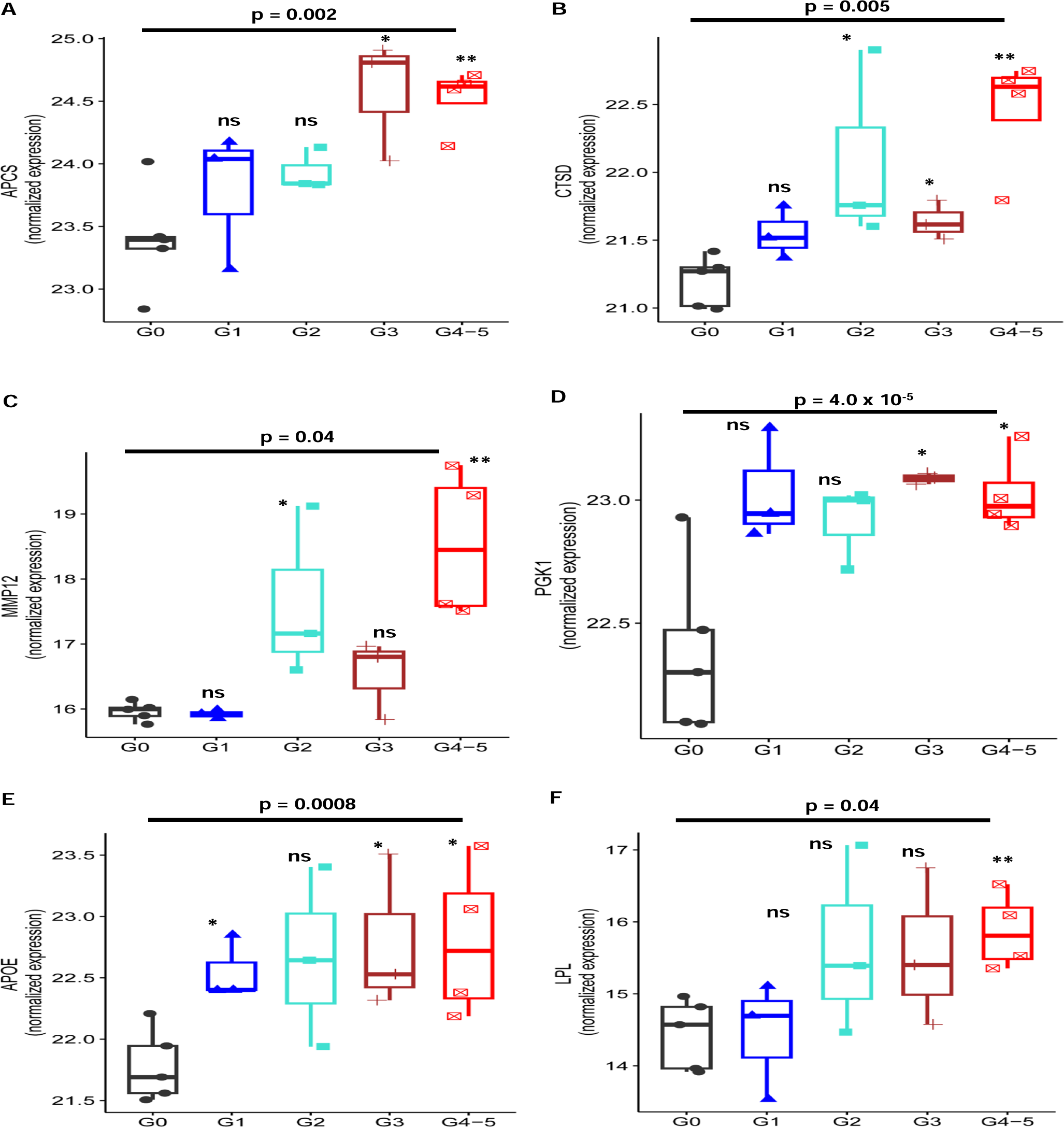
Boxplots summarizing the variation among the samples for representative proteins. A, Serum amyloid P-component (APCS); B, Cathepsin D (CTSD); C, Macrophage metalloelastase (MMP12); D, Phosphoglycerate kinase 1 (PGK1); E, Apolipoprotein E (APOE); and F, Lipoprotein lipase (LPL); Shown above the line are the nominal p values in case (G1 through G4-5 combined) vs. control (G0). Shown on the top of each of the boxes for the case groups (G1 through G4-5) represent the t test outcomes between each case group vs. the control (G0). The log2-transformed expression is shown on the y-axis. *, **, and ns stand for being significant at the 5%, 1% level, and not significant, respectively.

## Discussion

To our knowledge, this is the first study that aims to explore the proteomic profiling of the intracranial artery plaques from post-mortem human brains using mass spectrometry-based proteomic analysis. Despite of the small sample size, our study demonstrated that DEPs and GO terms are different in arterial segments with different degrees of atherosclerotic changes compared with those without atherosclerosis.

### DEP proteins are potential targets for biomarker discovery

Among the top 4 DEPs (LONP1, RPS19, MRPL12 and SNU13, **Fig. 2B**), none of them have been demonstrated to be associated with the atherosclerosis in the current literature. SNU13 is a protein involved in spliceosome assembly, its corresponding gene SNU13 belongs to the family of small nucleolar RNAs, which may contribute to the mechanism of atherosclerosis^44^. MRPL12 plays roles in mitochondrial translation. In one study, increased level of MRPL12 was found in left ventricle tissue of high-fat diet-fed mice with myocardial infarction (MI) than lose low-fat diet-fed mice with MI. In addition, MRPL12 level is also increased in atrial appendage tissue of diabetic patients with ischemic heart disease compared to non-diabetic patients. RPS proteins regulate multiple biological processes, such as chromatin structure, gene transcription, and RNA processing and splicing and post-translational modification^45^. RPS19, a component of the 40S ribosomal subunit, was found to interact with macrophage migration inhibitory factor (MIF) to attenuate its pro-inflammatory function of MIF^46^. In an animal study focusing on inflammatory kidney disease, RPS19 was found to be a potent anti-inflammatory agent, which appears to work primarily by inhibiting MIF signaling^47^. As a component of the mitochondrial large ribosomal subunit, LONP1 is an essential protease of the mitochondrial matrix and regulates crucial mitochondrial function including the degradation of oxidized proteins, folding of imported proteins and maintenance of the correct number of copies of mitochondrial DNA^48^. One study showed that upregulation of LONP1 could be beneficial as it reduces the cardiac injury by preventing oxidative damage of proteins and lipids, preserving mitochondrial redox balance and reprogramming bioenergetics by reducing Complex I content and activity^49^. The pathophysiological mechanism of the upregulation in these proteins in ICAD is unclear. We hypothesized that increased level of RPS19 and LONP1 may be protective effects in response to inflammatory process and oxidative damages, respectively, during the atherosclerosis genesis. The associations between these upregulated proteins and ICAD need to be further assessed, which may represent new research opportunities for discovery in future studies.

Among the top 50 proteins, several were found to be associated with cardiovascular or atherosclerotic disease. The adaptor protein FADD participates in the process of apoptosis.^50^ In a population-based cohort study, increased level of FADD was associated with higher incidence of coronary events during a mean follow-up of 19 years^35^. As a mitochondrial oxidoreductase, AIFM1 contributes to cell death programs and assembly of respiratory chain. Abnormal expression of AIFM1 leads to mitochondrial dysfunction which may contribute to mechanism of atherosclerosis as mitochondria plays an importance role in atherogenesis^51, 52^. In another study that performed serum proteomic study based on samples from “Munich Vascular Biobank”, AIFM1 was upregulated in patients with vulnerable carotid atherosclerotic lesions.^36^ In addition, immunostaining showed higher expression of AIFM1 in patients with unstable plaque, and enriched staining of AIFM1 was observed in regions where apoptosis occurred such as the necrotic core and the shoulder regions of the advanced CAP^36^. In one proteomic study based on the clot retrieved during acute thrombectomy from acute stroke patients (stroke etiology-predominantly atrial fibrillation), PGK1 was found to be positive in all the emboli and was associated with high LDL levels, however it does not appear to be specifically related to intracranial atherosclerosis and the study was underpowered to detected the PGK1 difference among different stroke etiologies given small sample size ^37^.

Previously identified proteins that were associated with extracranial atherosclerotic plaques, such as APCS, CTSD and MMP12 ^40–42^ were positively correlated with degree of the atherosclerotic changes in this study (**Figure 3A, B and C**), although they were not statistically significant DEPs after multiple test adjustment. This highlighted that the intracranial and extracranial atherosclerosis may have different proteomic signatures.

### DEP signatures are enriched for GO pathways relevant to ICAD

Among the top 20 most significantly upregulated pathways (**Figure 2C**), structural constituent of chromatin, nucleosome, protein -DNA complex, and DNA packing complex are the pathways involved in genetic activity such as transcription, packaging, rearrangement, replication and repair and posttranslational modifications; protein-lipid complex, plasma lipoprotein particle, and lipoprotein particle are pathways related to lipid metabolism; heparin binding, one type of glycosaminoglycan binding, is involved in vascular biology such as coagulation, anticoagulation and inflammation^53^. These findings are consistent with the previously reported pathways related to atherosclerosis. For example, chromatin remodeling was found to play important roles in cardiovascular disease^54^. In one study focusing on coronary artery disease, plasma nucleosome complexes were significantly elevated in patients with either severe coronary atherosclerosis or extremely calcified coronary arteries^55^. Consistent with current literature on genetic factors associated with intracranial atherosclerosis,^39^ our study showed the APOE and LPL gene were upregulated as the degree of the atherosclerosis worsene.

### Limitations

Firstly, one major limitation includes small sample size, as only 18 samples were included in the current study. Secondly, as some samples with different degree of atherosclerotic changes were from different subjects, along with the small sample size, it is challenging to adjust cofounders, such as demographics and cardiovascular risk factors. Lastly, the assessment of degree of atherosclerotic disease was based on visual assessment of fresh samples, which may not be as accurate as the assessment based on the fixed tissue; however it is not feasible to perform formalin fixation and hematoxylin and eosin staining on fresh tissue that will be prepared for proteomics, as the proteomic profiling will be affected by the formalin fixation process.

## Conclusions

Direct proteomic analysis of postmortem fresh-frozen intracranial artery samples by the MS-based proteomic technology is a feasible approach to identify proteins associated with atherosclerosis. Further proteomic profiling of plaques with a larger sample size is warranted to uncover mechanistic insights into ICAD and discover novel biomarkers that may help to improve early diagnosis, and predict disease progression & clinical outcome in ICAD.

## Ethics approval and consent to participate

All experimental procedures were conducted in accordance with the NIH guidelines for research. For the post-mortem samples that are used in the study, institutional review board (IRB) approval is not applicable per IRB committee at the Icahn School of Medicine at Mount Sinai (ISMMS).

## Informed Consent Statement

Not applicable.

## Institutional Review Board Statement

The study protocol was reviewed by institutional review board (IRB) at the Icahn School of Medicine at Mount Sinai (ISMMS), and the IRB determined that the proposed research is not research involving human subjects as defined by DHHS and FDA regulations. IRB review and approval by this organization is not required.

## Availability of data and materials

All codes are available up request.

## Authors’ contributions

Conceptualization, Q.H.; Investigation, E.W., Q.H., J.W., Z.P, J.P.; Resources, J.C., J.P., E.T., B.Z.; Writing – Original Draft, Q.H., E.W.; Writing – Review & Editing, All Authors; Supervision, J.P., B.Z.

## Acknowledgments

None.

---

**Figure S1.**
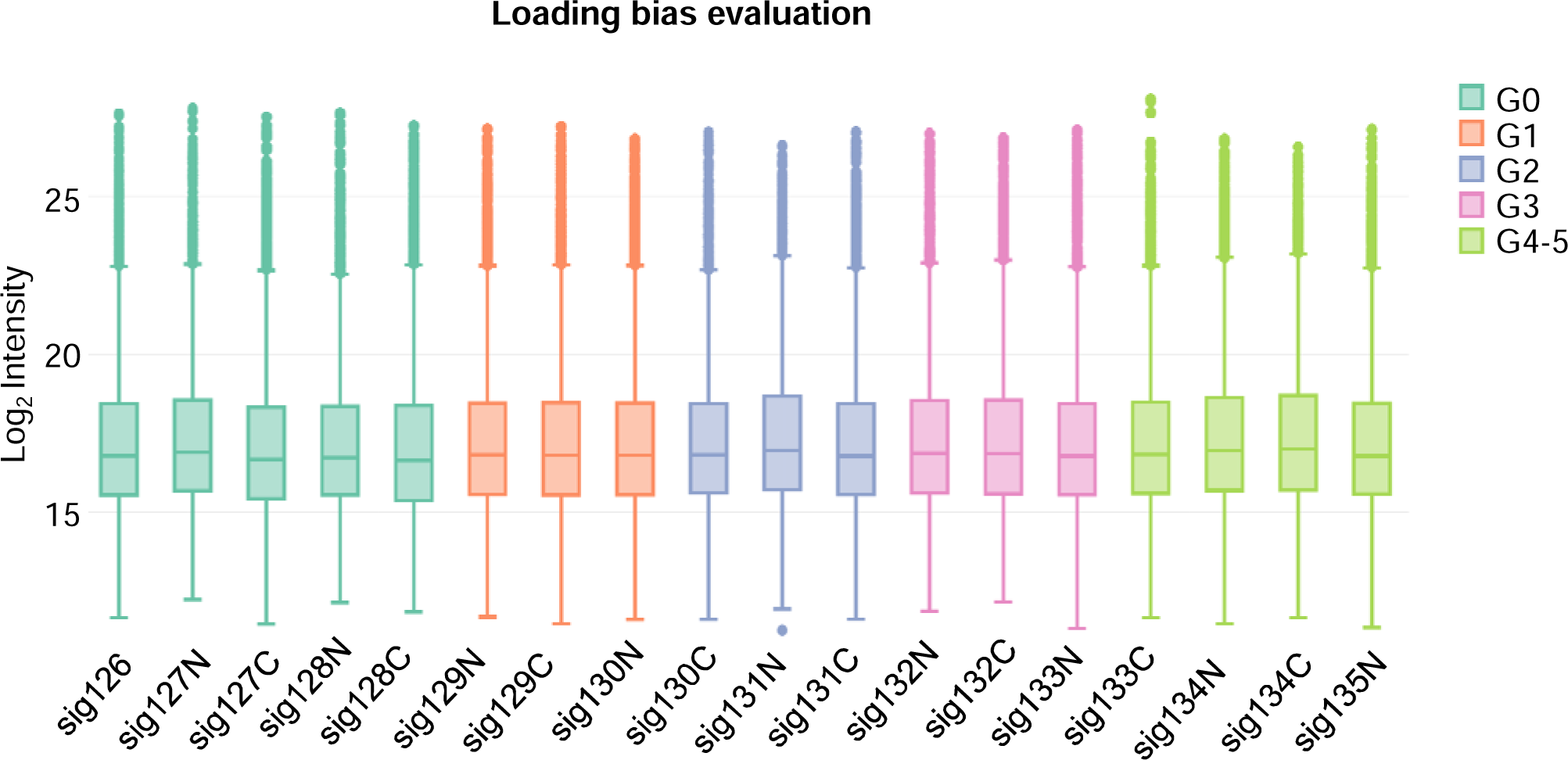
Boxplots showing the protein intensities across the samples. The results indicated no obvious loading bias occurred across the samples.

**Figure S2.**
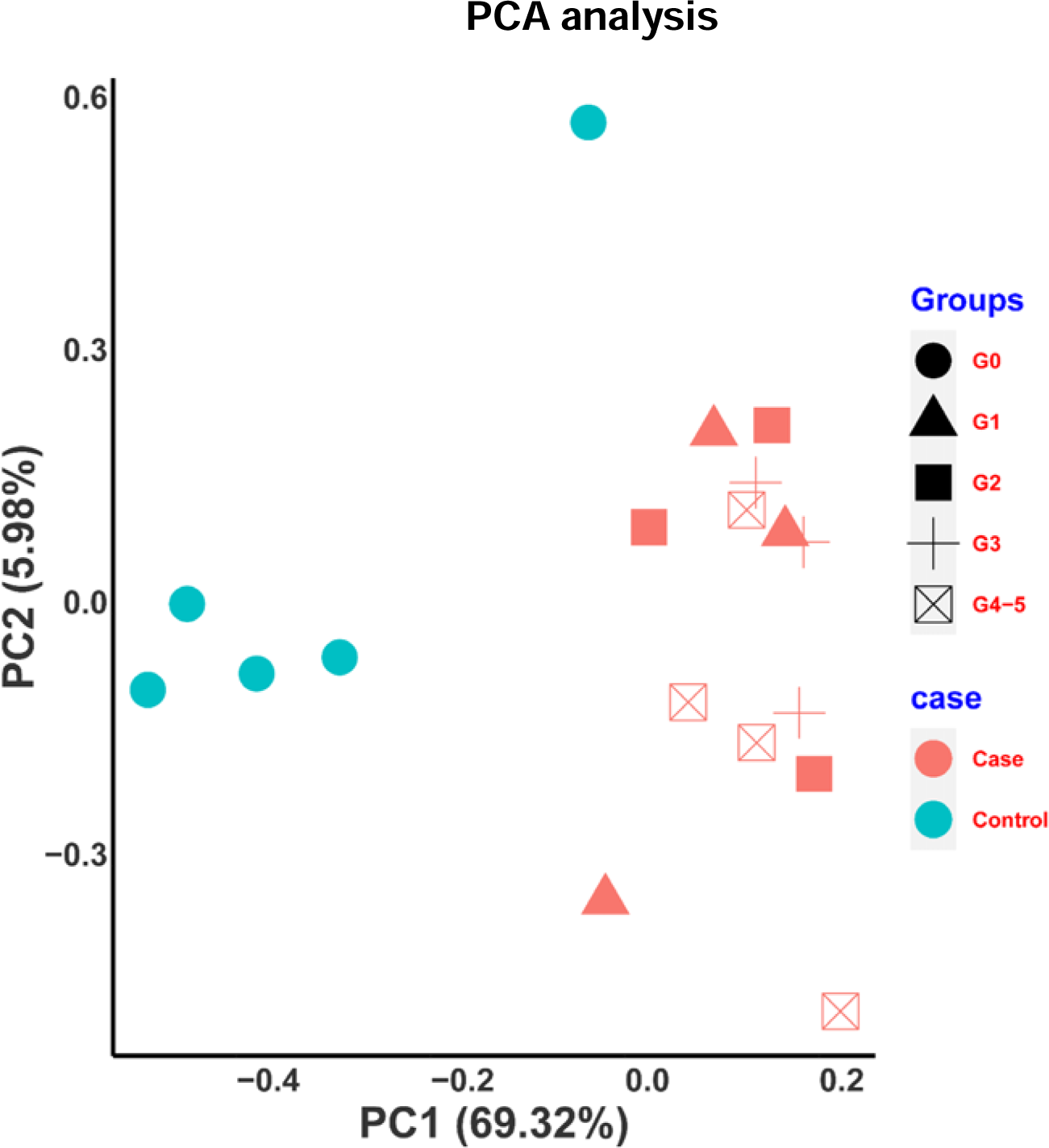
Principal Component analysis (PCA) of protein expression across the samples. Individual samples can be roughly grouped according to their clinical classification in case vs control.

## Supplementary data

Supple.data1.xlsx: Demographic and clinical traits of the samples

Supple.data2.xlsx: Whole proteome profiling of plaques in human brain samples by TMT-LC/LC-MS/MS

Supple.data3.xlsx: Differentially expressed proteins in case (G1 through G5) vs. control (G0). Supple.data4.xlsx: Gene ontology enrichment analysis of the DEPs.

## Reference

1. Sacco RL, Kargman DE, Gu Q, Zamanillo MC. Race-ethnicity and determinants of intracranial atherosclerotic cerebral infarction. The Northern Manhattan Stroke Study. Stroke; a journal of cerebral circulation 1995;26:14–20.

2. Wong LK. Global burden of intracranial atherosclerosis. Int J Stroke 2006;1:158–159.

3. Gorelick PB, Wong KS, Bae HJ, Pandey DK. Large artery intracranial occlusive disease: a large worldwide burden but a relatively neglected frontier. Stroke; a journal of cerebral circulation 2008;39:2396–2399.

4. Dearborn JL, Zhang Y, Qiao Y, et al. Intracranial atherosclerosis and dementia: The Atherosclerosis Risk in Communities (ARIC) Study. Neurology 2017;88:1556–1563.

5. Chimowitz MI, Lynn MJ, Derdeyn CP, et al. Stenting versus aggressive medical therapy for intracranial arterial stenosis. N Engl J Med 2011;365:993–1003.

6. Holmstedt CA, Turan TN, Chimowitz MI. Atherosclerotic intracranial arterial stenosis: risk factors, diagnosis, and treatment. Lancet Neurol 2013;12:1106–1114.

7. Bang OY. Intracranial atherosclerosis: current understanding and perspectives. J Stroke 2014;16:27–35.

8. Hurford R, Wolters FJ, Li L, Lau KK, Kuker W, Rothwell PM, Oxford Vascular Study Phenotyped C. Prevalence, predictors, and prognosis of symptomatic intracranial stenosis in patients with transient ischaemic attack or minor stroke: a population-based cohort study. Lancet Neurol 2020;19:413–421.

9. Verrills NM. Clinical proteomics: present and future prospects. Clin Biochem Rev 2006;27:99–116.

10. Chandramouli K, Qian PY. Proteomics: challenges, techniques and possibilities to overcome biological sample complexity. Hum Genomics Proteomics 2009;2009.

11. Geyer PE, Holdt LM, Teupser D, Mann M. Revisiting biomarker discovery by plasma proteomics. Mol Syst Biol 2017;13:942.

12. Puig N, Jimenez-Xarrie E, Camps-Renom P, Benitez S. Search for Reliable Circulating Biomarkers to Predict Carotid Plaque Vulnerability. Int J Mol Sci 2020;21.

13. Rifai N, Gillette MA, Carr SA. Protein biomarker discovery and validation: the long and uncertain path to clinical utility. Nat Biotechnol 2006;24:971–983.

14. Nakayasu ES, Gritsenko M, Piehowski PD, et al. Tutorial: best practices and considerations for mass-spectrometry-based protein biomarker discovery and validation. Nat Protoc 2021;16:3737–3760.

15. Bagnato C, Thumar J, Mayya V, et al. Proteomics analysis of human coronary atherosclerotic plaque: a feasibility study of direct tissue proteomics by liquid chromatography and tandem mass spectrometry. Mol Cell Proteomics 2007;6:1088–1102.

16. Porcelli B, Ciari I, Felici C, et al. Proteomic analysis of atherosclerotic plaque. Biomed Pharmacother 2010;64:369–372.

17. Baragetti A, Mattavelli E, Grigore L, Pellegatta F, Magni P, Catapano AL. Targeted Plasma Proteomics to Predict the Development of Carotid Plaques. Stroke; a journal of cerebral circulation 2022:101161STROKEAHA122038887.

18. Herrington DM, Mao C, Parker SJ, et al. Proteomic Architecture of Human Coronary and Aortic Atherosclerosis. Circulation 2018;137:2741–2756.

19. Zhou Y, Yuan J, Fan Y, et al. Proteomic landscape of human coronary artery atherosclerosis. Int J Mol Med 2020;46:371–383.

20. Nunez E, Fuster V, Gomez-Serrano M, et al. Unbiased plasma proteomics discovery of biomarkers for improved detection of subclinical atherosclerosis. EBioMedicine 2022;76:103874.

21. Rocchiccioli S, Pelosi G, Rosini S, et al. Secreted proteins from carotid endarterectomy: an untargeted approach to disclose molecular clues of plaque progression. J Transl Med 2013;11:260.

22. Nehme A, Kobeissy F, Zhao J, Zhu R, Feugier P, Mechref Y, Zibara K. Functional pathways associated with human carotid atheroma: a proteomics analysis. Hypertens Res 2019;42:362–373.

23. Liang W, Ward LJ, Karlsson H, Ljunggren SA, Li W, Lindahl M, Yuan XM. Distinctive proteomic profiles among different regions of human carotid plaques in men and women. Sci Rep 2016;6:26231.

24. Stakhneva EM, Striukova EV, Ragino YI. Proteomic Studies of Blood and Vascular Wall in Atherosclerosis. Int J Mol Sci 2021;22.

25. Lind L, Gigante B, Borne Y, et al. Plasma Protein Profile of Carotid Artery Atherosclerosis and Atherosclerotic Outcomes: Meta-Analyses and Mendelian Randomization Analyses. Arterioscler Thromb Vasc Biol 2021;41:1777–1788.

26. Nurmohamed NS, Belo Pereira JP, Hoogeveen RM, et al. Targeted proteomics improves cardiovascular risk prediction in secondary prevention. Eur Heart J 2022;43:1569–1577.

27. Ridker PM. Proteomics for the prediction and prevention of atherosclerotic disease. Eur Heart J 2022;43:1578–1581.

28. Bleijerveld OB, Zhang YN, Beldar S, Hoefer IE, Sze SK, Pasterkamp G, de Kleijn DP. Proteomics of plaques and novel sources of potential biomarkers for atherosclerosis. Proteomics Clin Appl 2013;7:490–503.

29. Barth RF, Kellough DA, Allenby P, Blower LE, Hammond SH, Allenby GM, Buja LM. Assessment of atherosclerotic luminal narrowing of coronary arteries based on morphometrically generated visual guides. Cardiovasc Pathol 2017;29:53–60.

30. Wang Z, Kavdia K, Dey KK, et al. High-throughput and Deep-proteome Profiling by 16-plex Tandem Mass Tag Labeling Coupled with Two-dimensional Chromatography and Mass Spectrometry. J Vis Exp 2020.

31. Wang X, Li Y, Wu Z, Wang H, Tan H, Peng J. JUMP: a tag-based database search tool for peptide identification with high sensitivity and accuracy. Mol Cell Proteomics 2014;13:3663–3673.

32. Ritchie ME, Phipson B, Wu D, Hu Y, Law CW, Shi W, Smyth GK. limma powers differential expression analyses for RNA-sequencing and microarray studies. Nucleic Acids Res 2015;43:e47.

33. Chen SY, Feng Z, Yi X. A general introduction to adjustment for multiple comparisons. J Thorac Dis 2017;9:1725–1729.

34. Yu G, Wang LG, Han Y, He QY. clusterProfiler: an R package for comparing biological themes among gene clusters. OMICS 2012;16:284–287.

35. Xue L, Borne Y, Mattisson IY, et al. FADD, Caspase-3, and Caspase-8 and Incidence of Coronary Events. Arterioscler Thromb Vasc Biol 2017;37:983–989.

36. Bischoff LP, J., Paloschi V., Maegdefessel, L. Abstract MP18: Mitochondrial Inducing Factor 1 Affects Stabilization Of Advanced Atherosclerotic Lesions In Carotid Artery Disease. Vascular Discovery: From Genes to Medicine Scientific Sessions Seattle, Washington, USA 2021.

37. Rao NM, Capri J, Cohn W, et al. Peptide Composition of Stroke Causing Emboli Correlate with Serum Markers of Atherosclerosis and Inflammation. Front Neurol 2017;8:427.

38. Abboud S, Viiri LE, Lutjohann D, et al. Associations of apolipoprotein E gene with ischemic stroke and intracranial atherosclerosis. Eur J Hum Genet 2008;16:955–960.

39. Liu M, Gutierrez J. Genetic Risk Factors of Intracranial Atherosclerosis. Curr Atheroscler Rep 2020;22:13.

40. Fasehee H, Fakhraee M, Davoudi S, Vali H, Faghihi S. Cancer biomarkers in atherosclerotic plaque: Evidenced from structural and proteomic analyses. Biochem Biophys Res Commun 2019;509:687–693.

41. Mahdessian H, Perisic Matic L, Lengquist M, et al. Integrative studies implicate matrix metalloproteinase-12 as a culprit gene for large-artery atherosclerotic stroke. J Intern Med 2017;282:429–444.

42. Malaud E, Merle D, Piquer D, et al. Local carotid atherosclerotic plaque proteins for the identification of circulating biomarkers in coronary patients. Atherosclerosis 2014;233:551–558.

43. Munshi A, Babu MS, Kaul S, Rajeshwar K, Balakrishna N, Jyothy A. Association of LPL gene variant and LDL, HDL, VLDL cholesterol and triglyceride levels with ischemic stroke and its subtypes. Journal of the neurological sciences 2012;318:51–54.

44. Farina FM, Weber C, Santovito D. The emerging landscape of non-conventional RNA functions in atherosclerosis. Atherosclerosis 2023;374:74–86.

45. Bhavsar RB, Makley LN, Tsonis PA. The other lives of ribosomal proteins. Hum Genomics 2010;4:327–344.

46. Filip AM, Klug J, Cayli S, et al. Ribosomal protein S19 interacts with macrophage migration inhibitory factor and attenuates its pro-inflammatory function. J Biol Chem 2009;284:7977–7985.

47. Lv J, Huang XR, Klug J, et al. Ribosomal protein S19 is a novel therapeutic agent in inflammatory kidney disease. Clin Sci (Lond) 2013;124:627–637.

48. Shin CS, Meng S, Garbis SD, et al. LONP1 and mtHSP70 cooperate to promote mitochondrial protein folding. Nat Commun 2021;12:265.

49. Venkatesh S, Li M, Saito T, et al. Mitochondrial LonP1 protects cardiomyocytes from ischemia/reperfusion injury in vivo. J Mol Cell Cardiol 2019;128:38–50.

50. Gupta S, Kim C, Yel L, Gollapudi S. A role of fas-associated death domain (FADD) in increased apoptosis in aged humans. J Clin Immunol 2004;24:24–29.

51. Bano D, Prehn JHM. Apoptosis-Inducing Factor (AIF) in Physiology and Disease: The Tale of a Repented Natural Born Killer. EBioMedicine 2018;30:29–37.

52. Madamanchi NR, Runge MS. Mitochondrial dysfunction in atherosclerosis. Circ Res 2007;100:460–473.

53. Munoz EM, Linhardt RJ. Heparin-binding domains in vascular biology. Arterioscler Thromb Vasc Biol 2004;24:1549–1557.

54. Rosa-Garrido M, Karbassi E, Monte E, Vondriska TM. Regulation of chromatin structure in the cardiovascular system. Circulation journal : official journal of the Japanese Circulation Society 2013;77:1389–1398.

55. Borissoff JI, Joosen IA, Versteylen MO, et al. Elevated levels of circulating DNA and chromatin are independently associated with severe coronary atherosclerosis and a prothrombotic state. Arterioscler Thromb Vasc Biol 2013;33:2032–2040.

